# Genetic modifiers of APOBEC-induced mutagenesis

**DOI:** 10.1101/2023.04.05.535598

**Authors:** Tony M. Mertz, Elizabeth Rice-Reynolds, Ly Nguyen, Anna Wood, Nicholas Bray, Debra Mitchell, Kirill Lobachev, Steven A. Roberts

**Affiliations:** School of Molecular Biosciences and Center for Reproductive Biology, Washington State University, Pullman, WA 99164, USA; School of Biological Sciences, Georgia Institute of Technology, Atlanta, GA 30332, USA

**Keywords:** APOBEC cytidine deaminases, mutagenesis, histone H3 K56 acetylation, CTF18-RFC complex, BRCA1 and BRCA2, breast and cervical cancer

## Abstract

The cytidine deaminases APOBEC3A and APOBEC3B (A3B) are prominent mutators of human cancer genomes. However, tumor-specific genetic modulators of APOBEC-induced mutagenesis are poorly defined. Here, we utilized a screen to identify 61 gene deletions that increase A3B-induced mutations in yeast. Also, we determined whether each deletion was epistatic with UNG1 loss, which indicated whether the encoded factors participate in the error-free bypass of A3B/Ung1-dependent abasic sites or suppress A3B-catalyzed deamination by protecting against aberrant formation of single stranded DNA (ssDNA). Additionally, we determined that the mutation spectra of A3B-induced mutations revealed genotype-specific patterns of strand-specific ssDNA formation and nucleotide incorporation across APOBEC-induced lesions. Combining these three metrics we were able to establish a multifactorial signature of APOBEC-induced mutations specific to (1) failure to remove H3K56 acetylation, which results in extremely high A3B-induced mutagenesis, (2) defective CTF18-RFC complex function, which results in high levels of A3B induced mutations specifically on the leading strand template that synergistically increase with loss of UNG1, and (3) defective HR-mediated bypass of APOBEC-induced lesions, which were epistatic with Ung1 loss and result from increased Rev1-mediated C-to-G substitutions. We extended these results by analyzing mutation data for human tumors and found BRCA1/2-deficient breast cancer tumors display 3- to 4-fold more APOBEC-induced mutations. Mirroring our results in yeast, for BRCA1/2 deficient tumors Rev1-mediated C-to-G substitutions are solely responsible for increased APOBEC-signature mutations and these mutations occur on the lagging strand during DNA replication. Together these results identify important factors that influence the dynamics of DNA replication and likely the abundance of APOBEC-induced mutation during tumor progression as well as a novel mechanistic role for BRCA1/2 during HR-dependent lesion bypass of APOBEC-induced lesions during cancer cell replication.

## Introduction

Tumorigenesis is driven by the occurrence of mutations in oncogenes and tumor suppressor genes. Additional mutations allow tumors to grow, metastasize, and develop resistance to chemotherapeutics. Recently, large-scale whole genome sequencing of tumors has identified 81 mutation signatures as indexed by the Catalogue of Somatic Mutations in Cancer (COSMIC) (Alexandrov et al. 2020) that describe the processes mediating tumor genome evolution. Although many mutation signatures have been attributed to endogenous and exogenous sources of DNA damage, defects in DNA repair, or decreased DNA replication fidelity (Lynch and de la Chapelle 2003; Imai and Yamamoto 2008; Cancer Genome Atlas 2012; Cancer Genome Atlas Research et al. 2013; Palles et al. 2013; Alexandrov et al. 2020), the mechanisms that generate many mutation signatures are either poorly understood or unknown.

Two of the most prominent mutation signatures, SBS2 and SBS13, in COSMIC’s list of mutation signatures (Alexandrov et al. 2020), are primarily comprised of C-to-T or C-to-G mutations, respectively, within TCW trinucleotide motifs (with W being either an A or T). These two mutation signatures are second only to aging-signature mutations in their contribution to mutation burden in cancer (Alexandrov et al. 2013), are over-represented in 15% of all sequenced tumors (Alexandrov et al. 2013) and in many tumors constitute more than 50% of all mutations (Burns et al. 2013b; Roberts et al. 2013). These mutation signatures have been attributed to APOBEC deaminases, which normally play diverse biological roles in the innate and adaptive immune responses (Refsland and Harris 2013). When dysregulated, APOBEC family members mutate genomic DNA by converting deoxycytidine to deoxyuridine (dU) within regions of single-stranded DNA (ssDNA) most commonly during DNA replication, but also during transcription and DNA repair (Chaudhuri et al. 2003; Nik-Zainal et al. 2012; Roberts et al. 2012; Lada et al. 2013; Kazanov et al. 2015; Haradhvala et al. 2016; Hoopes et al. 2016). Nuclear localization (Bogerd et al. 2006; Lackey et al. 2013), a preference for deaminating cytidine in TC or TCW motifs *in vitro* (Harris et al. 2002; Harris et al. 2003; Bishop et al. 2004; Yu et al. 2004; Dang et al. 2006; Henry et al. 2009; Burns et al. 2013a; Adolph et al. 2017) and TCW motifs within genomic DNA of cells (Hoopes et al. 2016), detectible *ex cellular* deamination activity (Cortez et al. 2019), positive correlation between expression and the load of signature mutations in tumors (Burns et al. 2013a; Burns et al. 2013b; Roberts et al. 2013), and reduced APOBEC signature mutations in cultured human cells with CRISPR mediated APOBEC3A (A3A) knockouts (Petljak et al. 2022) all point to APOBEC3 family members A3A and A3B being responsible for the large number of C-to-T or C-to-G mutations within TCW motifs found in cancer genomes.

APOBEC activity causes both primary driver mutations and additional late sub-clonal driver mutations in cancer (de Bruin et al. 2014; Henderson et al. 2014; McGranahan et al. 2015; Cancer Genome Atlas Research et al. 2017; Jamal-Hanjani et al. 2017). In addition, abundance of APOBEC signature mutations and APOBEC expression have prognostic value in respect to patient outcomes and response to cancer therapeutics (Sieuwerts et al. 2014; Walker et al. 2015; Billingsley et al. 2016; D’Antonio et al. 2016; Law et al. 2016; Middlebrooks et al. 2016; Yan et al. 2016; Glaser et al. 2018; Van Hoeck et al. 2019; Alexandrov et al. 2020). Due to these strong links to cancer progression and survival, several studies have characterized defects that reduce cell viability when combined with APOBEC expression (Mehta et al. 2020; Biayna et al. 2021), However, the modulators of APOBEC-induced mutagenesis in tumor cells are largely unknown. Messenger RNA expression of A3A and A3B correlate with the number of APOBEC-induced mutations in tumors, implicating transcriptional dysregulation as one contributing factor. However, the correlation is generally weak, which suggests that additional factors such as protein stability, post-translational modifications, substrate availability, and repair of APOBEC-induced dU play significant roles in modulating APOBEC-induced mutagenesis in tumors.

To identify genetic factors that modulate APOBEC-induced mutagenesis, we generated and screened a collection of *Saccharomyces cerevisiae* deletion strains expressing A3B for increased APOBEC-dependent mutations. We measured *CAN1* mutation rates for all the screen hits and a group of candidates with functions related to the identified hits and discovered 61 gene deletions that increase A3B-induced mutagenesis. We produced *CAN1* mutation spectra for nearly all the deletion strains with increased A3B-induced mutagenesis to confirm that the mutations were APOBEC-dependent and to determine how these defects effect stand bias and repair choice of APOBEC-induced dU. Also, we combined the identified deletions with a deletion of *UNG1,* which is required for both C-to-G mutation-generating translesion synthesis and error-free lesion bypass of dU-dependent abasic sites and measured A3B-dependent mutation rates. Analysis of these results allowed us to place the genes that most appreciably protect against increased A3B mutagenesis into four main groups, those that encode (1) proteins that participate in DNA replication, replication fork stability, and restart, which likely increase ssDNA availability, (2) proteins that allow for homologous recombination (HR)-dependent error-free bypass of APOBEC-dependent abasic sites, (3) Ctf18-RFC complex members, which function in sister chromatid cohesion and PCNA loading, and (4) chromatin modifiers and remodelers. The large number of genes involved in recombination prompted us to determine if defective recombinational repair affects APOBEC-induced mutagenesis in tumors. We found that tumors with loss of BRCA1/BRCA2 function are enriched for APOBEC mutagenesis specifically for C-to-G mutations in TCW motifs, which indicates that HR-dependent error-free bypass of abasic sites generated by APOBEC and UNG2 limits mutagenesis in human tumors.

## Results

In tumor cells, multiple factors likely combine to determine the number of mutations that result from the activity of APOBEC cytidine deaminases. Due to the difficulty of using mutagenesis in human cell lines as a phenotype for a genetic screen, we utilized the budding yeast *S. cerevisiae* as a model system to screen for genes preventing APOBEC-induced mutagenesis. We created a library of haploid yeast deletion strains expressing A3B using a mating, sporulation, and selection strategy (Figure 1A). We measured mutation frequencies in three successive rounds, which allowed us to identify ORF deletions (screen hits) that increased the frequency of canavanine resistant (Can^R^) colonies at least 2-fold. This set of hits contained most of the gene deletions previously identified as modulators of increased APOBEC-induced mutations including *ung1*Δ, *mph1*Δ, and *tof1*Δ (Hoopes et al. 2016; Hoopes et al. 2017), supporting the effectiveness of the screen strategy. Based on the known functions and interactors of the screen hits, we also created a list of additional candidate genes with probable roles in restricting APOBEC-induced mutagenesis (see Supplemental Table 1). To validate whether all candidates truly increased A3B-induced mutation, we transformed either haploid strains from the BY4741 yeast deletion library (i.e., not created via the mating/sporulation method), or *de novo* deletion strains constructed from the wild-type BY4741 yeast with either an

**Figure 1:**
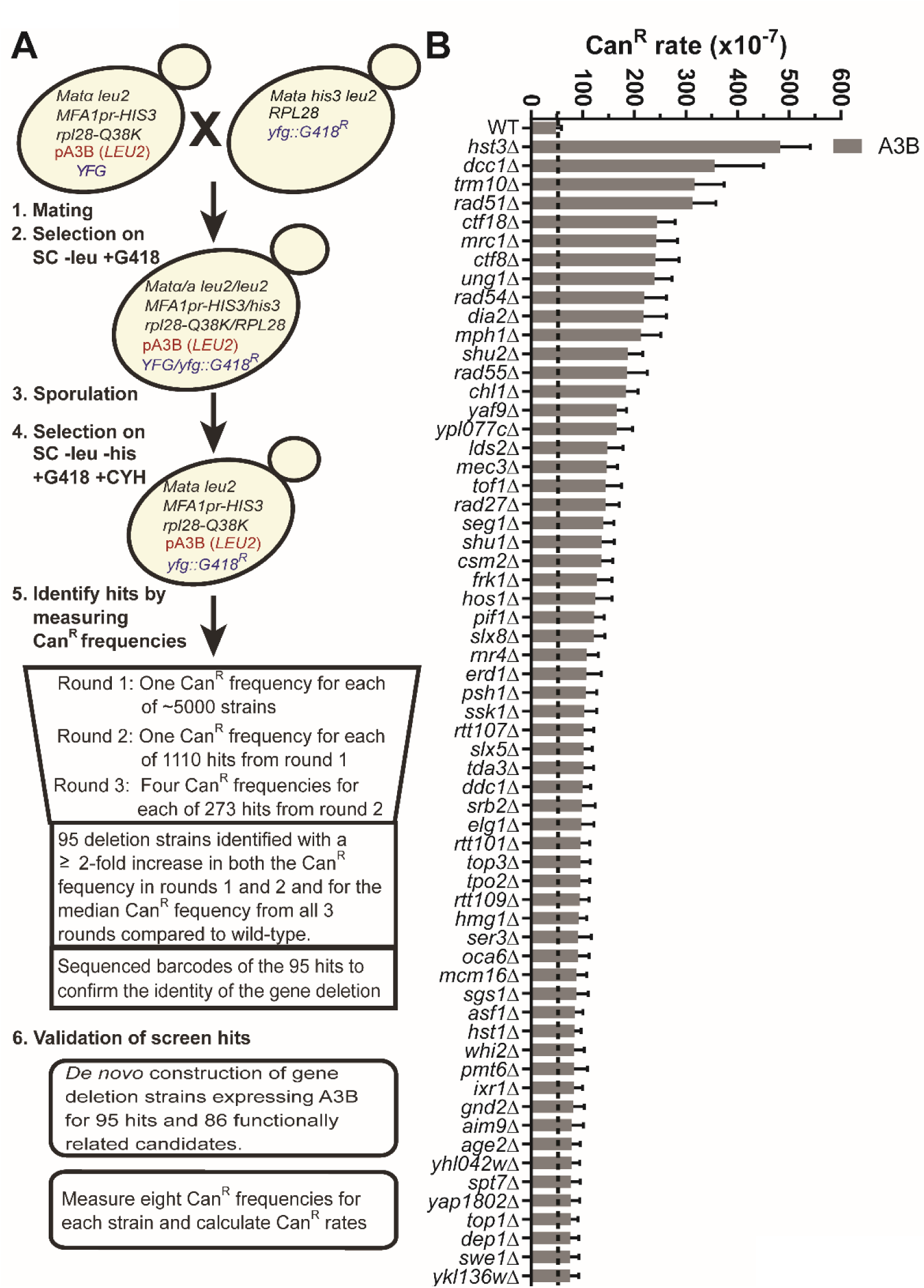
Identification of yeast gene deletions that increase APOBEC3B-induced mutations. (A) Schematic of the yeast screen used to identify genes that elevate A3B-induced mutagenesis. Haploid yeast containing a single gene deletion and the A3B expression plasmid were obtained by mating *MATα* yeast carrying the A3B expression plasmid with each *MATa* strain of the yeast gene deletion library, sporulating the resulting diploid cells and selecting the desired haploid genotype with appropriate selective media The resulting haploid cells were subjected to serial rounds of Can^R^ frequency measurements to identify gene deletions that likely augment A3B-induced mutagenesis. To confirm these deletions resulted in increased A3B-induced mutations we re-created each deletion strain expressing A3B de novo and measured Can^R^ mutation rates B) Can^R^ rates for 61 yeast gene deletions that elevate A3B-induced mutation greater than 1.55-fold over the A3B-induced Can^R^ rate in wild-type yeast (grey dashed line) without overlapping 95% confidence intervals and 1.55-fold higher than additive for spontaneous deletion-dependent Can^R^ and wild-type with A3B Can^R^ rates Error bars indicate 95% confidence intervals. For all mutation rate data see Supplemental Table 2.

A3B expression plasmid and separately an empty vector control, and measured *CAN1* mutation rates (Supplemental Table 2). Gene deletions present in yeast strains that met the following metrics were classified as *bona fide* modulators of APOBEC-induced mutagenesis: (1) 95% confidence intervals for *CAN1* mutation rates were non-overlapping for the wild-type strain expressing A3B and deletion strain expressing A3B (2) A3B-induced mutation rate was >1.55-fold higher in deletion strain compared to wild-type, (3) A3B-induced mutation rate was >1.55-fold higher than the sum of the spontaneous Can^R^ rate (for that strain) and the rate for the wild-type strain with A3B expression, which ensures the increase in the mutation rate is due to elevated A3B-induced mutagenesis and not an additive effect for strains with a relatively high spontaneous mutation rate (Supplemental Table 2). In total, we identified 61 gene deletions that increase A3B-induced mutagenesis (Figure 1B). Deletion of *HST3*, members of the Ctf18-RFC complex, *TRM10*, *MRC1*, HR factors, and *UNG1* were among those with the greatest increase in A3B-induced mutation.

Gene ontology (GO) analysis of the 61 validated defects that increased A3B-induced mutagenesis indicated this set was highly enriched for genes with functions in DNA intra-S and replication checkpoints, histone deacetylation, DNA replication, sister chromatid cohesion, DNA repair, recombinational repair, and chromatid segregation (Figure 2A). Analysis of interactions between these genes via String-DB.org indicate that homologous recombination, histone H3 lysine 56 acetylation (H3K56Ac), and the CTF18-RFC complex likely play significant roles in APOBEC-induced mutagenesis (Figure 2B). However, numerous interactions between genes of different clusters (Figure 2B) and our observation that many genes within this data set are associated with multiple complexes and/or functional processes (Figure 2C) made it difficult to determine which function(s) of individual genes act to limit APOBEC-generated mutations from mutation rates alone.

**Figure 2:**
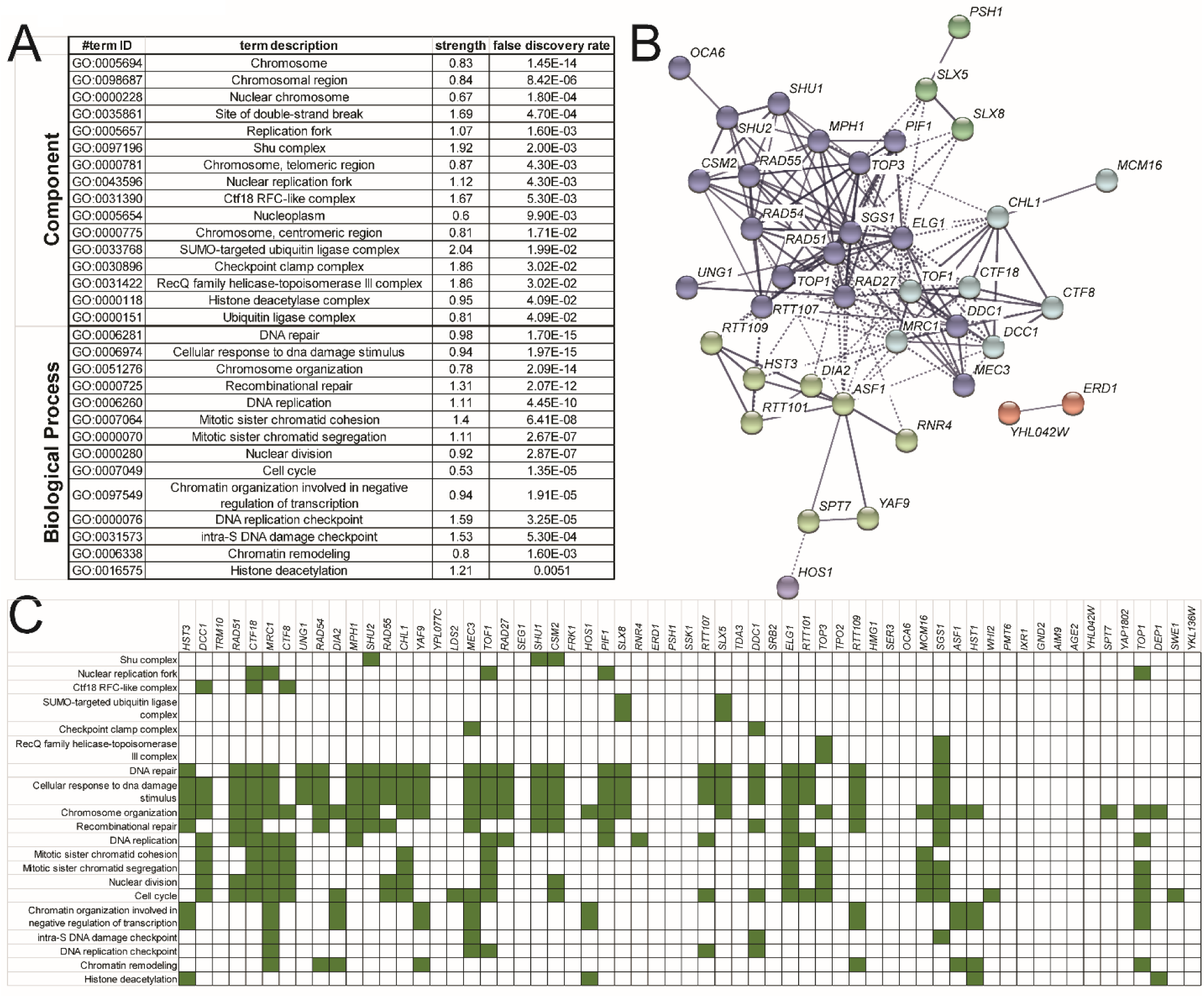
Molecular processes enriched among gene deletions that increase A3B-induced mutagenesis. (A) Gene ontology analysis for molecular processes that are enriched among gene deletions that increase A3B-induced mutation. Only categories with strengths equal to or greater than one and having at least three genes represented in the category are shown. Categories for similar processes are represented with the specific GO process with the lowest FDR-corrected p-value. (B) Functional association network for proteins that limit A3B-induced mutation as created by String-DB.org. The 61 gene deletion strains that were confirmed to increase A3B-induced mutagenesis were used in creating the network. Highest confidence linkage settings and MCL clustering with inflation parameter of three were employed. Proteins without connections to other proteins in the network are hidden. Each MCL clustering node is indicated by a unique color. (C) Gene deletions that elevate A3B-induced mutation function in multiple GO processes. The specific enriched GO processes that each gene contributes to are indicated in green.

To determine if the genes identified reduce APOBEC-induced mutations by functioning in repair of APOBEC-generated dU or the resulting abasic sites, we measured mutation rates in deletion strains also lacking *UNG1*. In yeast, repair of APOBEC-induced dUs and prevention of mutations requires the activity of a uracil DNA glycosylase, Ung1 *(Hoopes et al. 2016)*, and in *ung1*Δ strains all APOBEC-induced dU result in C-to-T mutations, because deoxyuridine templates like deoxythymidine. Therefore, deletions combined with *ung1*Δ that significantly increase the APOBEC-induced *CAN1* mutation rates above *ung1*Δ alone likely expand the availability of ssDNA that serves as substrate for APOBEC activity. In contrast, deletions that are epistatic to *ung1*Δ for *CAN1* mutation rates in the presence of A3B likely function during error-free bypass of abasic sites created by combined activities of A3B and Ung1. Therefore, we expressed A3B in yeast strains containing combined deletions that increased APOBEC-induced mutagenesis and *ung1*Δ and measured mutation rates (Figure 3). Supporting previous findings, we found that deletion of *MPH1* (Hoopes et al. 2017) and Shu-complex members *CSM2*, *SHU1*, and *SHU2* (Rosenbaum et al. 2019), which were previously shown to be important for error-free lesion bypass (Godin et al. 2016), were epistatic to *ung1*Δ for A3B-induced mutagenesis. We also found epistasis between genes encoding proteins with direct roles in HR (i.e., *RAD51*, *RAD54*, and *RAD55*) and recombination-intermediate resolution (i.e., *SGS1* and *TOP3) (Fabre et al. 2002)*. Deletion of *MEC3* and *DDC1*, which are members of the yeast 9-1-1 complex, were also epistatic with *ung1*Δ supporting previous resulting suggesting that they have a role in error-free lesion bypass (Karras et al. 2013). Conversely, deletion of components of replication checkpoint surveillance complex, *MRC1* and *TOF1,* synergistically increased the rate of A3B-induced mutagenesis when combined with *ung1*Δ, consistent with their roles at stalled replication forks (Katou et al. 2003; Tourriere et al. 2005; Hodgson et al. 2007). Interestingly, deletion of *CTF18* and *CTF8*, which are members of the CTF18-RCF complex that has roles as an alternate PCNA loader and unloader (Bylund and Burgers 2005), in sister chromatid cohesion (Mayer et al. 2001), in HR (Ogiwara et al. 2007), and in checkpoint activation (Garcia-Rodriguez et al. 2015), also synergistically increased the rate of A3B-induced mutagenesis when combined with *ung1*Δ, suggesting one, or more, of these function(s) reduce levels of ssDNA.

**Figure 3:**
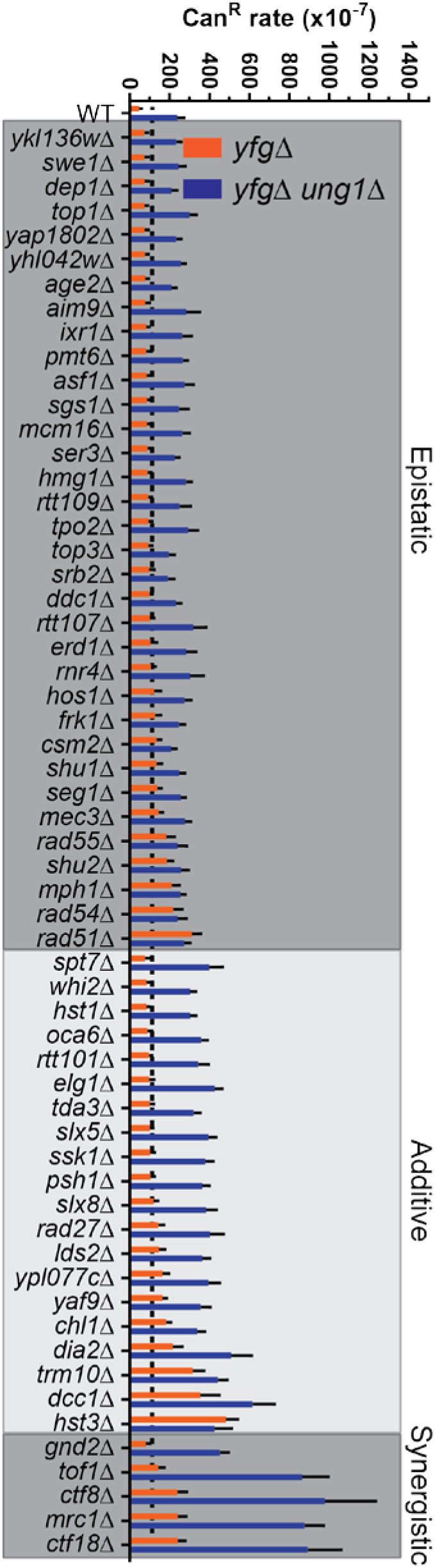
Impact of ung1Δ in mutants exhibiting increased A3B-induced mutation. A3B-induced Can^R^ rates in yeast that contain genes that limit A3B-induced mutation *(yfg*Δ: orange bars) or in yeast with genes that limit A3B-induced mutation co-deleted with *UNG1 (yfg*Δ *ung1*Δ, blue bars). Error bars indicated 95% confidence intervals. Double deletion strains *(yfg*Δ *ung1*Δ) that maintain similar A3B-induced Can^R^ rates to *ungi*Δ strains (black dashed line) are considered epistatic with *ungi*Δ and likely contribute to the repair or error-free bypass of A3B-induced lesions, *yfg*Δ *ung1*Δ strains with A3B-induced Can^R^ rates higher than *ungi*Δ strains are either additive or synergistic with *UNG1* deletion and likely increase the amount of ssDNA that A3B acts upon.

To gain additional insight as to how individual gene deletions increase A3B-induced mutagenesis, we sequenced *CAN1* mutants from each deletion strains expressing A3B. Consistent with our conclusion that the identified genes protect against APOBEC-induced mutations, greater that 70% of mutations for each genotype expressing A3B involved C-to-G or C-to-T substitutions at TC dinucleotides (83% for all genotypes combined) and therefore were attributable to A3B activity (Supplemental Tables 3 and 4, Figure 4). We and others have shown that APOBECs primary target the leading strand template during DNA replication (Bhagwat et al. 2016; Haradhvala et al. 2016; Hoopes et al. 2016; Morganella et al. 2016; Seplyarskiy et al. 2016; Saini et al. 2017; Sui et al. 2020). However, there are multiple means by which modulators of APOBEC-induced mutagenesis could affect the mutational strand bias. In wild-type BY4741 strain with A3B expression, the lagging strand template is associated with 2.1-fold more APOBEC-induced mutations for the *CAN1* coding sequence (C nucleotides are favored over G nucleotides at *CAN1* in this genomic location). Like *ung1*Δ, many deletions of genes with likely roles in HR-dependent error-free bypass (*sgs1*Δ, *rad54*Δ, *rad55*Δ, *csm2*Δ, and *mph1*Δ) had a significant increase in mutations at C nucleotides, while others (*shu2*Δ and *rad51*Δ) had a similar, but not statistically significant increase (Figure 4A), indicating loss of error-free template switching emphasizes the lagging strand template bias of A3B-induced mutation. Interestingly, defects with the largest increase in lagging strand bias include *dia2*Δ, *seg1*Δ, *erd1*Δ, *yaf9*Δ, and *ypl077c*Δ lack established roles in error-free lesion bypass. Conversely, defects in all three members of the CTF18-RCF complex, *ctf18*Δ, *ctf8*Δ, and *dcc*1Δ (Figure 4A) resulted in a leading strand template mutation bias, which indicated increased ssDNA on the leading strand template.

**Figure 4:**
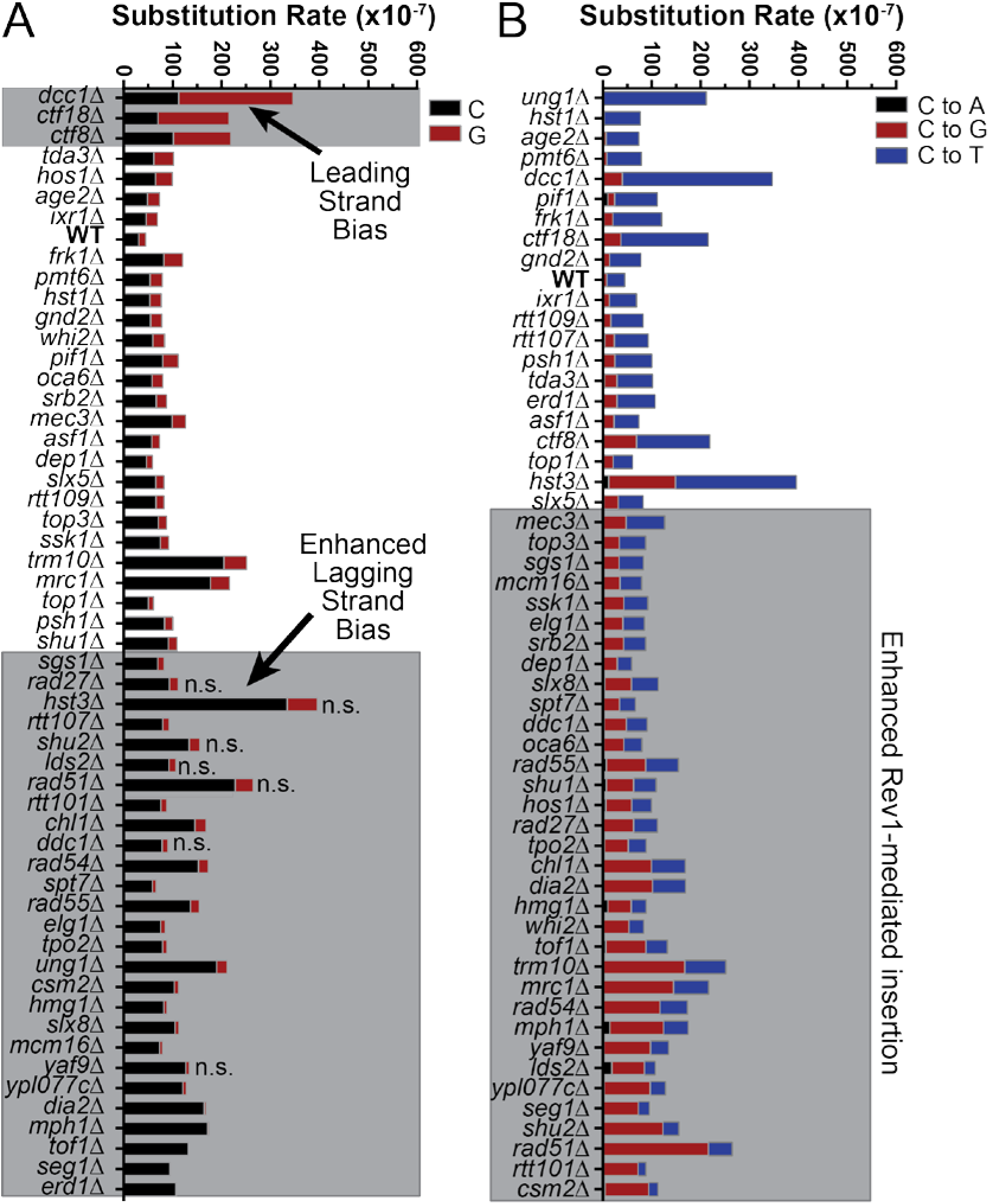
Spectra of A3B-induced mutations in *CAN1.* The *CAN1* gene of independent Can^R^ isolates was amplified and sequenced to determine the spectra of A3B-induced mutations in yeast with specific gene deletions. Note that A3B-induced *CAN1* mutants were not sequenced from *aim9*Δ*, rnr4*Δ*, ser3*Δ*, swel*Δ*, yap1802*Δ*, YKL136W*Δ, and *YHL042W*Δ strains. Combined, 83% of mutations involved C or G substitutions at TC sites, indicative of A3B activity. (A) Strand bias of A3B-induced mutations in *CAN1* is influenced by specific gene deletions. The rate of C (black bars) or G (red bars) substitutions in each genotype of yeast expressing A3B. Wild-type yeast expressing A3B favors C substitutions over G substitutions indicating that the lagging strand template is the top DNA strand of *CAN1* at its location on ChrV. The ratio of A3B-induced C-to-G substitutions in wild-type yeast were compared to the same ratio for each yeast deletion strain by two-sided Fisher’s Exact Test. Genotypes with p-values < 0.00083 (based on Bonferroni multiple hypothesis testing correction) were considered to have altered strand bias. (B) The rates of C-to-A, C-to-T, and C-to-G substitutions (complementary substitutions were combined) in *CAN1* for yeast expressing A3B and containing with various gene deletions. The ratio of C-to-T and C-to-G substitutions were compared pairwise between wild-type yeast expressing A3B and each specific gene deletion strain by two-sided Fisher’s Exact Test. Gene deletion strains displaying p-values < 0.00083 (for Bonferroni correction) were deemed to have greater usage of Rev1 catalytic activity during bypass of A3B-induced abasic sites.

Analysis of the *CAN1* mutation spectra also allowed us to determine how gene deletions affected the ratio of C-to-G to C-to-T A3B-induced mutations with higher ratios being indicative of increased REV1-mediated TLS to bypass A3B-dependent abasic sites. In contrast, lower C- to-G to C-to-T ratios indicate either enhanced A-rule bypass or decreased Ung1 activity (Figure 4B). As expected, in *ung1*Δ stains, only A3B-induced C-to-T substitutions were observed.

Deletions causing defects in HR-dependent error-free bypass significantly increased the C-to-G to C-to-T ratio and the increase of the *CAN1* mutation rate in these strains is almost entirely due to increased C-to-G mutations. In addition, elevated A3B-mutagenesis in *dia2*Δ, *mrc1*Δ, and *tof1*Δ stains was primarily driven by increased C-to-G mutations, indicating that factors maintaining replication fork stability also decrease TLS bypass that uses REV1-deoxycytidyl transferase activity. Deletion of factors influencing H3 K56 acetylation (i.e., *HST3, RTT107, ASF1)* and disruption of the CTF18-RCF complex (*ctf18*Δ, *ctf8*Δ, or *dcc*1Δ), however, increased both the rate of C-to-G and C-to-T substitution.

We next utilized unsupervised hierarchical clustering of gene deletion strains based upon changes in strand bias, substitution pattern, epistatic relationship with *ung1*Δ, and Can^R^ rate, to see if these features stemmed from general disruption of the common pathways or specific functions of individual genes. As expected, distinct nodes where observed based on this clustering, with one containing all members of the CTF18-RCF complex (*ctf18*Δ, *ctf8*Δ, or *dcc*1Δ), another being associated with replication fork stability factors (*mcm16*Δ, *dia2*Δ, and *tof1*Δ), and a third containing a large number of factors involved in homologous recombination (*top3a*Δ, *rad55*Δ, *rad54*Δ, *rad51*Δ, *shu1*Δ, *shu2*Δ, and *csm2*Δ) (Figure 5, data from Supplemental Table 9). This indicates that the mutagenic features of the genes in these nodes (e.g., leading strand bias for CTF18-RFC factors, epistasis of recombination factors with *ung1*Δ, and increased Rev1 utilization in recombination deficient strains) are generalizable to the entire pathway. However, some notable deviations included clustering of *mph1*Δ with replication fork stability genes and *mrc1*Δ clustering in an undefined node and indicate that some gene specific functions may also influence these metrics.

**Figure 5:**
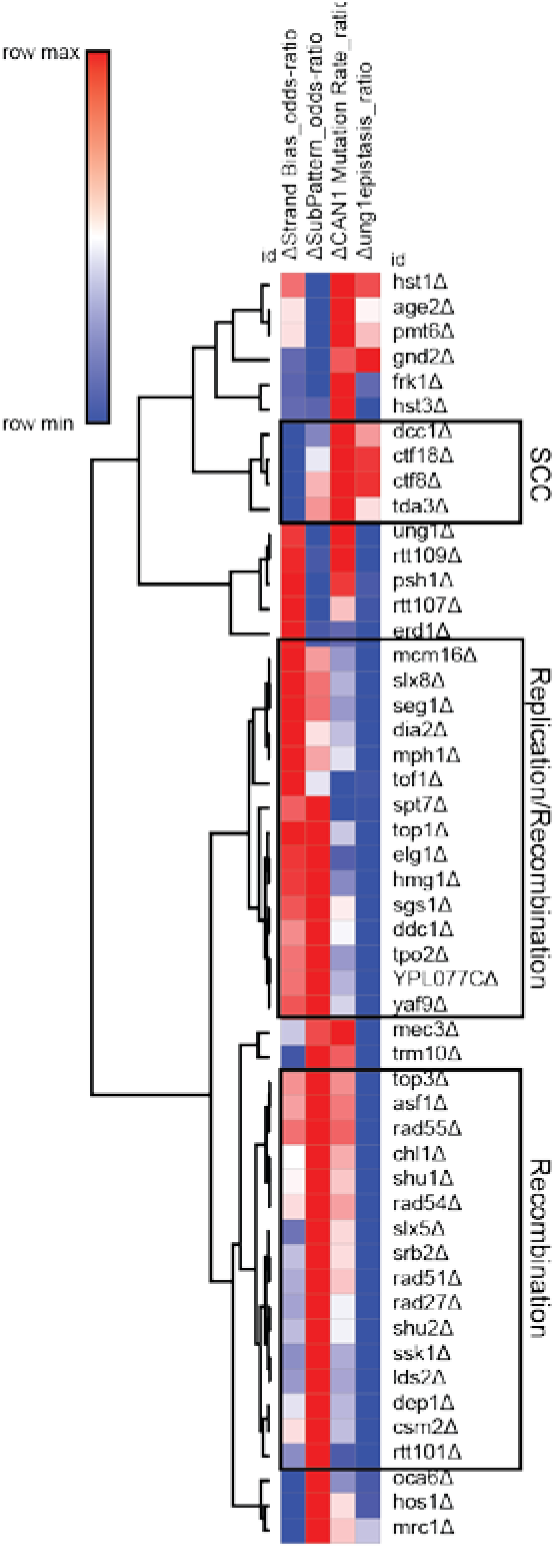
Non-biased hierarchical clustering of yeast gene deletions that similarly impact A3B-induced mutation. Yeast gene deletion strains for which greater than 20 independent A3B-induced *CAN1* mutations were sequenced were clustered using average linkage based upon (1) how much the gene deletion elevated A3B-induced mutation over that in wild-type yeast, ACAN1 Mutation Rate_ratio, which equals the log2(Can^R^ rate_vfgΔ_/Can^R^ rate_wild-type_), (2) how much the rate of mutation in the yfgΔ *ung1*Δ co-deletion strains compared to the mutation rate in *ung1*Δ *A* strains, Δung1epistasis_ratio, which equals the (log2(Can^R^ rate_yfgΔ *ung1*Δ_/Can^R^ rate*_ung1_*_Δ_), (3) mutational strand bias, ΔStrand Bias_odds-ratio, which equals the Iog2(substitutions at C/substitutions at G nucleotides), and (4) polymerase usage, ΔSubPattern_odds-ratio, which equals the log2(C-to-T substitutions/C-to-G substitutions). Nodes of genes associated with sister chromatin cohesion (SCC), DN A replication and recombination, and recombination were identified (black boxes).

Upon finding that multiple proteins involved in HR limit APOBEC3B-induced mutation in yeast, we assessed whether orthologous factors in human tumors have a similar role. We obtained somatic mutations from 560 whole-genome sequenced breast cancers and assessed the relative abundance of SBS2 (i.e., TCW to TTW) and SBS13 (i.e., TCW to TGW) in each tumor. Among these tumors, 61 originated from individuals with germline mutations in either *BRCA1* or *BRCA2*, which are required for HR-repair activities in human cells (Figure 6A). We stratified these tumors into either BRCA1/2-proficient or BRCA1/2-deficient classes and determined the median abundance of total APOBEC signature mutations (i.e., SBS2 and SBS13 combined), SBS2 mutations, and SBS13 mutations alone in each class. We found that that BRCA1/2-deficient tumors contained a 2.4-fold higher median load of total APOBEC-induced mutations (p<0.0001 by Mann Whitney test) compared to BRCA1/2-proficient tumors (Figure 6B). Surprisingly, this increase is largely due to a 3.8-fold elevation in SBS13 mutations (p<0.0001 by Mann Whitney test), while SBS2 mutations occurred at similar levels between BRCA1/2-proficient and deficient tumors (median values of 151 and 142 SBS2 mutations per tumor), respectively. The increase of only SBS13 mutations in BRCA1/2-deficient breast cancers mirror our observation of increased APOBEC-induced mutation are driven by higher rates of C-to-G substitutions in recombination-deficient yeast strains (Figure 4B). Therefore, this result suggests that BRCA1 and BRCA2 likely limit APOBEC signature mutations in human cancer cells by participating in HR-mediated error-free lesion bypass of base lesions in the lagging strand template. Consequently, APOBEC-induced mutations in recombination-defective yeast maintain a replicative asymmetry, which indicates most mutations originate from A3A or A3B deaminating the cytidines in the lagging strand template. To determine whether BRCA1 and BRCA2 likely facilitate error-free lesion bypass of APOBEC-induced DNA damage in breast cancer tumors, we assessed whether SBS13 mutations have replicative asymmetry associated with deamination of the lagging strand template in the absence of BRCA1 or BRCA2 (Figure 6C). As previously reported for pan-cancer analysis of APOBEC-induced mutations, SBS13 mutations in BRCA1/2-proficient breast cancers followed the direction of replication, associating strongly with deamination of cytidines in the lagging strand template. SBS13 mutations in BRCA1/2-deficient tumors also displayed the same replicative asymmetry, indicating the primary source of APOBEC-induced mutation in the tumors was due to deamination ssDNA at the replication fork. This result supports a model where the increase in APOBEC-induced mutation in these tumors likely results from defective error-free lesion bypass as opposed increased ssDNA that could occur in BRCA1/2-deficient tumors.

**Figure 6:**
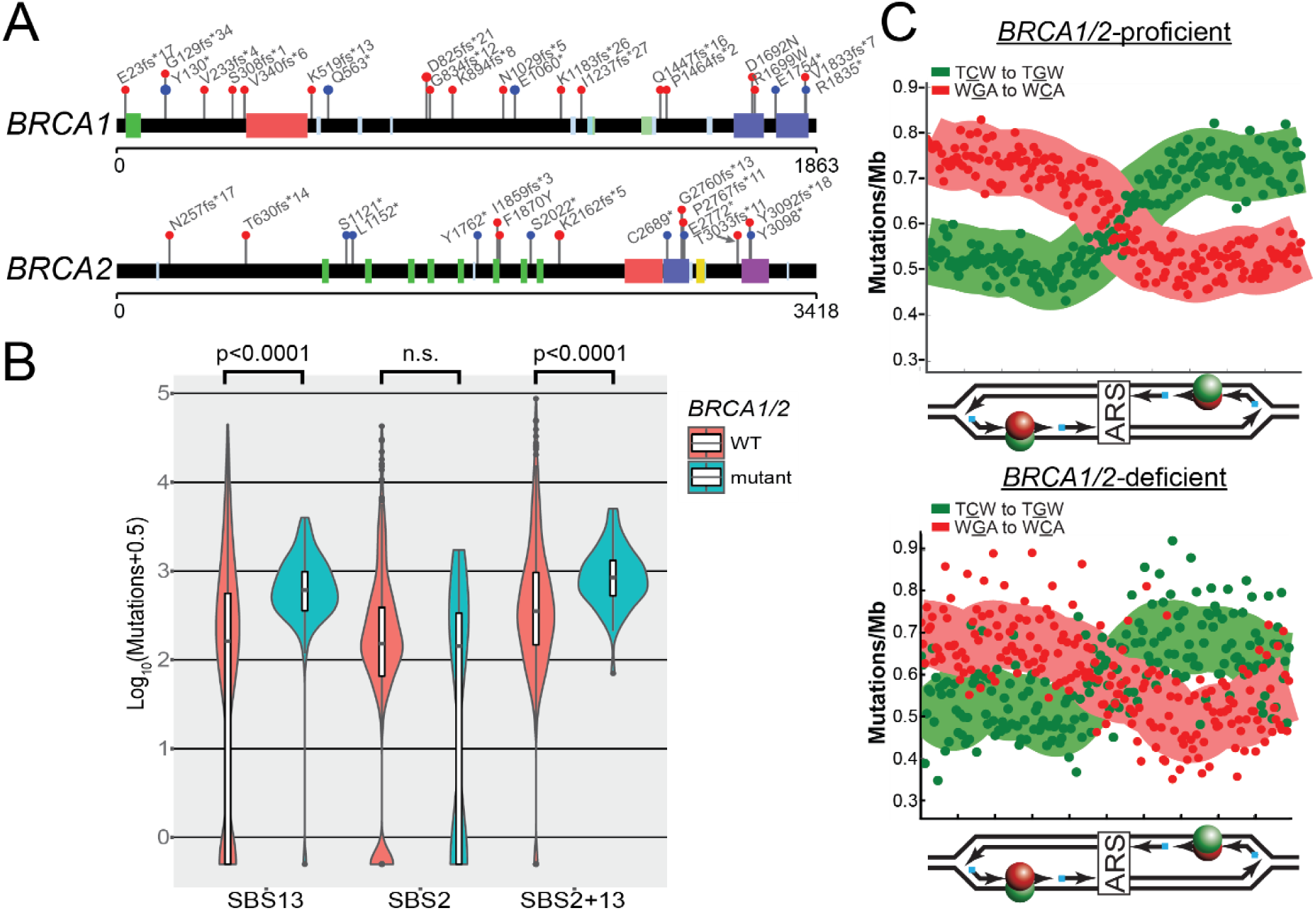
Recombination deficient breast cancers have elevated APOBEC signature mutagenesis. (A) Position of germline mutations in *BRCA1* and *BRCA2* present in 560 breast cancers sequences as part of the International Cancer Genome Consortium. (B) Median estimated number of COSMIC SBS2, SBS13, and SBS2+SBS13 mutations in breast cancers from BRCA-proficient and BRCA-mutant patients. (C) Replication strand bias of SBS13 mutations in tumors from BRCA-proficient and BRCA-deficient patients.

We next assessed whether a reduction of other recombination factors have effects on the abundance of APOBEC signature mutations like those observed in BRCA1/2 deficient tumors. We obtained somatic base substitutions, copy number variations, and RNA-seq expression data from 309 CESC tumors characterized by the Cancer Genome Atlas (TCGA). Among these tumors, we identified 148 samples that contained a statistical over-representation of APOBEC signature mutations. By GISTIC analysis, a single copy of either *RAD51*, *RAD51C*, or *HPRT1* were lost in 31, 12, and 30 of the APOBEC-mutated CESC tumors, respectively. Tumors containing single *RAD51* and *RAD51C* alleles displayed 1.42-fold and 1.16-fold lower median gene expression than copy neutral tumors (p=0.0002 and p=0.0314 respectively by Mann Whitney test) (Figure 7A), indicating that the loss of one allele of these genes corresponds to a lower transcript level and likely decreases protein abundance. Consequently, a single allele deletion of either *RAD51* or *RAD51C*, but not the housekeeping gene *HPRT1*, in tumors correlated with higher minimal estimates of total APOBEC-induced mutations (p=0.015 and p=0.0243, respectively by Mann Whitney Test) (Figure 7B). Importantly, within this CESC dataset, the overall number of copy number changes within a tumor does not correlate with the number of base substitutions (Figure 7C), ensuring that the associations we observed between somatic single gene deletion in *RAD51* and *RAD51C* are unlikely to result from tumors with higher copy number burden, and thus more likely to have deletions in *RAD51* or *RAD51C,* having de facto higher mutation burdens. Thus, single allele deletions of *RAD51* and *RAD51C* appear to be haplo-insufficient in limiting APOBEC-induced mutation and support a general role for HR factors in error-free bypassing of APOBEC-induced DNA damage in multiple human cancer types.

**Figure 7:**
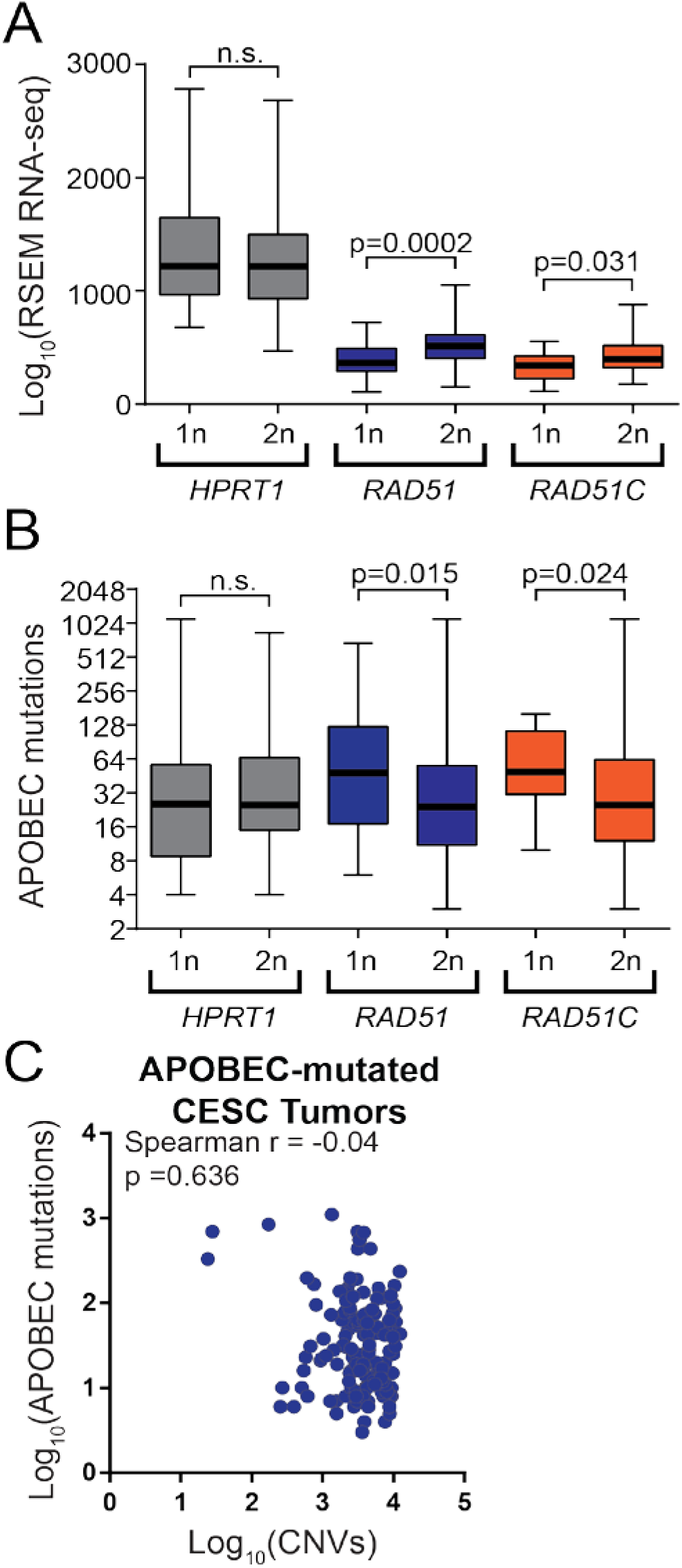
Impact of recombination gene haplo-insufficiency on APOBEC-induced mutagenesis in cervical cancers. (A) Median RSEM normalized RNA-seq expression values (middle horizontal bar in each box plot) for *HPRT1, RAD51,* and *RAD51C* in cervical cancers 1 or 2 copies of each gene as determined by GISTIC analysis of SNP6 array data. Error bars indicate the range of RSEM expression values among tumors in each group. Tumors with two copies of *RAD51* and *RAD51C* have higher expression of the respective genes than tumors with one copy as assessed by Mann-Whitney ranked sum test. (B) Minimal estimate of the number of APOBEC-induced mutations in cervical cancers with 1 or 2 copies of *HPRT1, RAD51*, and *RAD51C.* Median values are indicated by the middle bar in the box plots, while error bars indicate the range of values in each group. (C) Lack of correlation between the number of copy number variants and APOBEC-induced mutations in cervical cancers sequenced by The Cancer Genome Atlas. Spearman coefficient (rho) and p-value are indicated.

## Discussion

Utilizing a yeast deletion screen, we identified 61 strains with gene deletions that exhibited significantly increased A3B-induced mutagenesis (Figure 1B). Genes encoding proteins functioning in DNA replication, homologous recombination, replication, and chromatin modification were enriched among genes that protect against APOBEC-generated mutations (Figure 2). By measuring the A3B-induced mutation rate yeast strains containing each gene deletion combined with *ung1*Δ and generating *CAN1* mutation spectra from A3B-expressing strains containing single gene deletions, we were able to generate discrete characteristics associated with defective processes that increase APOBEC3B-induced mutagenesis (summarized in Figure 5).

We found that many gene deletions in yeast with primary and supporting roles in HR (i.e., *top3a*Δ, *rad55*Δ, *rad54*Δ, *rad51*Δ, *shu1*Δ, *shu2*Δ, *csm2*Δ, *sgs1*Δ, *mph1*Δ, *mec3*Δ, and *ddc1*Δ) increased APOBEC-induced mutagenesis. Almost all genes with roles in HR when deleted showed clear epistasis for the rate of A3B-induced *CAN1* mutation when combined with *ung1*Δ, which indicates they function downstream of Ung1 in error-free repair. Also, deletion of genes with roles in HR significantly increased the C-to-G to C-to-T ratio among sequenced *CAN1* mutants, which indicates greater usage of Rev1-mediated error-prone lesion bypass of APOBEC-dependent abasic sites and that the encoded proteins reduce APOBEC-induced mutations by participating in HR-dependent error-free lesion bypass (summarized in Figure 5). Additionally, we found that BRAC1/2 deficient tumors have an elevated abundance of APOBEC-induced mutations due solely to increases SBS13 mutations, which arise from REV1-dependent TLS (Petljak et al. 2022). These APOBEC-signature mutations in BRAC1/2 deficient tumors, have replication symmetry indicative of mutations arising from the activity of APOBECs on the lagging strand template during DNA replication. Although several studies indicate HR-dependent error-free bypass occurs in human cells (Adar et al. 2009; Taglialatela et al. 2021), this process remains poorly understood. Our results clearly indicate that BRCA1/2 function in an HR-dependent error-free bypass of lesions that block DNA replication, and that this repair pathway is used to reduce APOBEC-induced mutations in tumor cells.

The deletion of *HST3* resulted in the largest increase in the rate of A3B-induced *CAN1* mutations. Hst3 is a H3 histone deacetylase (HDAC) and member of the sirtuin family of deacetylases (Brachmann et al. 1995). Newly synthesized histones are acetylated by the actions of histone chaperone Asf1 and the histone acetyltransferase Rtt109 (Driscoll et al. 2007; Tsubota et al. 2007) to mark new histones that are incorporated into DNA behind the replication fork with H3 lysine 56 acetylation (H3K56Ac). Hst3 functions to deacetylate H3K56 during G2/M phases of the cell cycle. *hst3*Δ, *asf1*Δ, and *rtt109*Δ yeast strains elevated the rate A3B-induced mutagenesis at *CAN1* by 10.0-, 1.96-, and 1.77-fold respectively, which indicates both establishment and removal of H3K56Ac is important for preventing APOBEC-induced mutagenesis. Epistasis between *asf1*Δ and *rtt109*Δ and *ung1*Δ for A3B-induced mutagenesis likely results from the importance of HR on post-replicative establishment of H3K56ac (Munoz-Galvan et al. 2013). The A3B-induced mutation rate in *hst3*Δ stains is 1.8-fold higher than that of *ung1*Δ and strains with defects in error-free lesion bypass, which indicates *hst3*Δ increases the amount of ssDNA that can be deaminated by APOBECs. Combining *hst3*Δ with *asf1*Δ or *rtt109*Δ suppresses A3B-mutagenesis to levels observed in *asf1*Δ or *rtt109*Δ strains (Supplemental Table 2), which suggests persistent H3K56Ac is responsible for elevated A3B-induced mutagenesis in *hst3*Δ strains. This result is similar to those for spontaneous increases in the *CAN1* mutations rate for *hst3*Δ, *hst4*Δ strains which are repressed by loss of *ASF1* or *RTT109 (Kadyrova et al. 2013)*. Because defective H3K56Ac deacetylation has been implicated in control of origin firing, DNA replication efficiency, limiting break induced replication and mutations arising from replication, DNA damage response signaling, cohesion establishment, and HR (Thaminy et al. 2007; Celic et al. 2008; Munoz-Galvan et al. 2013; Che et al. 2015; Irene et al. 2016; Simoneau et al. 2016; Gershon and Kupiec 2021) further work is required to parse out the underlying mechanism(s) by which H3K56Ac influences APOBEC-induced mutation. Two additional deletions of genes encoding HDACs, *hos1*Δ and *hst1*Δ, also increased A3B-induced mutagenesis, but were dissimilar to *hst3*Δ for characteristics of A3B-induced mutagenesis (Figure 5). Deletions of other HDACs, *hda1*Δ, *hda2*Δ, *hos2*Δ, *hos3*Δ, *hst2*Δ, *hst4*Δ, and *sir2*Δ, did not significantly increase A3B-induced mutagenesis (Supplemental Table 2). In human cells, SIRT1, SIRT2, SIRT6, and HDAC1/2 have all been implicated in H3K56Ac deacetylation in different contexts (Das et al. 2009; Michishita et al. 2009; Yuan et al. 2009; Zhu et al. 2015). Investigating H3K56ac-mediated modulation of APOBEC-induced mutagenesis in tumors may be of significant importance given reports of elevated H3K56Ac in human tumor samples compared to normal tissue (Das et al. 2009), the potential of activated Ras-PI3K signaling to reduce H3K56Ac (Liu et al. 2012), and a large number of clinical trials for HDAC inhibitors that likely increase H3K56Ac levels in tumor cells (Eckschlager et al. 2017), all of which effect patient prognosis and influence therapeutic outcomes by increasing APOBEC-induced mutagenesis and for tumors with A3A or A3B expression.

Deletion of *CTF18*, *CTF8*, and *DCC1* caused some of the largest increases of APOBEC-induced mutagenesis we observed. The encoded proteins Ctf18, Cft8, and Dcc1 form the CTF18-RFC complex along with Rfc2. APOBEC-induced mutations within *CAN1* for *ctf8*Δ, *ctf18*Δ, and *dcc1*Δ were strongly biased to the leading strand and when these deletions were combined with *ung1*Δ, it led to a synergistic increase in the A3B-induced *CAN1* mutation rate, which indicates these deletions elevate APOBEC-generated mutagenesis by increasing the availability of ssDNA on the leading strand during DNA replication (summarized in Figure 5, and Supplemental Figure 1). Although CTF18-RFC has well characterized roles in formation of chromatid cohesion, characteristics of A3B-induced mutations in *ctf18*Δ, *ctf8*Δ, and *dcc1*Δ strains are different than other deleted genes with roles in cohesion (i.e., *sgs1*Δ, *top3*Δ, *chl1*Δ, and *mrc1*Δ). CTF18-RFC has been implicated in playing a role during HR (Ogiwara et al. 2007), however defective HR-dependent error-free lesion bypass tends to be epistatic with *ung1* and increase APOBEC-mutagenesis primarily on the lagging strand, which is contrary to the effect of defective CTF18-RFC upon APOBEC mutagenesis. Furthermore, CTF18-RFC was shown to participate in Mrc1-dependent activation Rad52 within the DNA replication checkpoint (Garcia-Rodriguez et al. 2015). However, defects in CTF18-RCF and *mrc1*Δ have opposing effects on both the strand bias and use of Rev1-mediated TLS in respect to APOBEC-induced mutations for *CAN1*, which indicates they affect APOBEC-induced mutagenesis in distinct ways. Taken together our results support studies that indicate but did not conclusively demonstrate, that PCNA loading by the CTF18-RFC complex is important the efficiency of replication and/or facilitating bypass DNA damage on the leading strand (Okimoto et al. 2016; Liu et al. 2020), which likely reduces ssDNA available for APOBEC activity on the leading DNA strand. This establishes novel outcomes of defective PCNA loading by CTF18-RFC, which have eluded previous studies.

In addition to genes with well characterized roles in DNA replication and repair, we identified many genes without clear roles that protect against A3B-mediated mutagenesis. Among those with the highest APOBEC-induced mutagenesis include *trm10*Δ, *dia2*Δ, *yaf9*Δ, and *ypl077c*Δ. *TRM10* encodes a tRNA methyltransferase without a known function in DNA metabolism, which does not have an obvious explanation for its modulation of A3B-induced mutagenesis. TRM10 is elevated upon induction of replication stress (Tkach et al. 2012), indicating it may be needed for recovery of these conditions, which could increase A3B-induced mutation. However, *TRM10* is adjacent to *RFC4*, and *trm10*Δ might increase replication stress by reducing *RFC4* expression. Deletions of *DIA2*, *YAF9*, and *YPL0177*c all increase lagging strand bias and Rev1 usage for A3B-induced mutations, which is consistent with the encoded proteins participating in HR-mediated bypass. The Yaf9-containing NuA4 histone deacetylase complex and Dia2 have been implicated in regulating HR (Bennett and Peterson 2015; Chaudhury and Koepp 2017), which suggests Yaf9 and Dia2 may participate in error-free bypass of APOBEC-induced abasic sites. However, when *DIA2*, *YAF9*, and *YPL0177*c deletions are combined with *ung1*Δ, they cause higher mutation rates than *ung1*Δ alone, which indicates these defects may also increase ssDNA. Consequently, these genes may protect against increasing APOBEC-induced mutagenesis via multiple functions and should be further investigated for additional roles in DNA repair and replication. We anticipate that additional genes not represented in the yeast genome, likely limit APOBEC-induced mutations in human cells. Our screen was limited to non-essential gene deletions, and it is likely that in tumor cells could also have elevated APOBEC-induced mutagenesis due to mutations, reduced expression, or altered post translational modifications that reduce the function of essential genes, like RPA, DNA polymerases (Sui et al. 2020) or other regulators of cell cycle progression.

Our finding that defective HR in BRCA and CESC tumors increases APOBEC-induced mutagenesis is a good indicator that many of the other genes we identified in yeast as restrictors of APOBEC mutagenesis may do so in the context of human tumors. Because APOBEC-induced dU produce well-defined, strand-specific abasic sites in the presence of UNG1/2, characterizing the spectra and distribution of APOBEC-induced mutations have the potential to further define functions of DNA repair and replication proteins with roles in bypass of abasic sites and those that protect against aberrant ssDNA formation in both yeast and human cells. For these reasons, we believe that in addition to identifying and characterizing novel modulators of APOBEC-induced mutagenesis, these finding will facilitate future efforts to understand the role of APOBECs in cancer biology.

## Methods

### Yeast growth conditions and strains

Yeast strains were grown on YPDA or synthetic complete (SC) dropout media (Burke et al. 2000). The strains used in this study are all derived from BY4742 or BY4741 and the library of BY4741 deletion strains was purchased from Open Biosystems (currently available from Transomic Technologies; Huntsville, AL). The genotype of strain HMK246, which was created by the Lobachev lab (Zhang et al. 2012) from BY4742, is: MATα, ura3-Δ, leu2-Δ, his3-Δ, lys2-Δ, rpl28-Q38K, mfa1Δ::MFA1pr-HIS3, V34205::lys2::(GAA)230, V29617::hphMX. Strain HMK246 was further modified by exchanging the hphMX selection cassette for natMX to create strain yTM-01. Strain yTM-01 was modified by converting the V34205::lys2::(GAA)230 allele to a functional *LYS2* allele to strain yTM-02. Strain yTM-03 was created by using a PCR generated cassette to delete *UNG1* in the yTM-02 strain. Strains used to screen for elevated A3B-induced mutagenesis were constructed as follows. First, strain yTM-02 was transformed with a *LEU2* marked ARS/CEN plasmid expressing APOBEC3B, pTM-021 (Rosenbaum et al. 2019). yTM-02+pTM21 was mated to each BY4741 deletion strain and diploids were selected on SC-leu+G418 media. The resulting diploid strains were replica plated to sporulation media. After 10 days, sporulation plates were replica plated to SC-his-leu+G418+cyclohexamide to select for haploid cells with individual gene deletions and expressing A3B. Clonal isolates of each deletion strain expressing A3B were generated by streaking to single colonies on SC-his-leu+G418. *UNG1* deletion strains were created by either transforming yeast strains with PCR generated NatMX deletion cassettes or by a mating yTM-03 and strains from the deletion library, sporulating, and selecting haploid yeast on SC-Leu+G418. For genotypes and additional details about individual strains, (see Supplemental Table 5) and for primer sequences used to construct and verify gene deletions (see Supplemental Table 6).

### Measurement of mutation frequencies and rates

To determine Can^R^ frequencies for yeast strains, colonies grown to ∼1 x 10^7^ cells were resuspended in H_2_O and ∼200 yeast cells were plated non-selective media and 2.5 x 10^5^ cells for A3B expressing strains, or 1 x 10^7^ cells for strains with the empty vector control were plated on SC-arg+canavanine media. *CAN1* mutation rates (i.e. mutations per generation) were calculated from ≥ 8 mutation frequency measurements by the maximum likelihood method (Zheng 2002) using either FALCOR (Hall et al. 2009) or calculated using the fluxer.R script (https://github.com/barricklab/barricklab/blob/master/fluxxer.R).

### Screen for genetic defects that increase APOBEC-induced mutations

The screening process for gene deletions that increase A3B-induced mutations was conducted in three rounds. In round one, a single mutation frequency was measured for each deletion strain expressing A3B. For deletion strains from round one with a Can^R^ frequency greater than or equal to 2-fold higher than the median mutation frequency, a second Can^R^ frequency was measured. For the 273 strains from round two with a Can^R^ frequency ≥2-fold higher than the median mutation frequency, new clonal isolates from the population of haploid cells were generated and colonies for Can^R^ frequencies were all grown to 4 x 10^6^ cells, and four additional Can^R^ frequencies were measured. deletion strains with a median mutation frequency ≥2-fold higher than that of wild-type BY4741 were labeled as hits from the screening process. The gene deletion in each of the 101 screen-derived candidates was verified by sequencing the unique barcode associated with deletion.

### Post screen characterization of screen hits and candidates

For the 101 screen-identified candidates and an additional 76 candidates that were chosen from GO-analysis of the screen-derived candidates, we obtained the original deletion strain from the BY4741 library or made the strain *de novo* from the parental BY4741 strain. These strains were transformed with either pySR-419, an empty vector control plasmid, or pSR-440, a hygromycinB resistance marked ARS/CEN plasmid expressing APOBEC3B (Hoopes et al. 2016) and used for mutation rate measurements and generation on mutation spectra.

### Determination of *CAN1* mutation spectra

Independent clonal yeast colonies were grown to ≈1 x 10^7^ cells and replica plated to SC-arg+canavanine media. Can^R^ papillae were subjected to an additional round of clonal expansion via single colony streaking. Genomic DNA was extracted from the clonal Can^R^ mutants using previously described techniques and used as templates for PCR amplification of *CAN1* utilizing primer pairs with unique combinations of barcodes, (Supplemental Table 6). Approximately 2500 amplicons were subjected to Single Molecule, Real-Time (SMRT) sequencing on either a PacBio RSII or PacBio Sequel instruments.

### Data Availability

Somatic mutations from whole genome sequenced breast cancers from the International Genome Consortium were obtained from https://dcc.icgc.org/api/v1/download?fn=/release_26/Projects/BRCA-EU/simple_somatic_mutation.open.BRCA-EU.tsv.gz. Presence of germline mutations in the *BRCA1* and *BRCA2* genes were listed in Supplementary Table 20 in (Nik-Zainal et al. 2016). Abundances of Signature 2 and Signature 13 mutations per tumor were obtained from Supplementary Table 21 in (Nik-Zainal et al. 2016). Copy number variations, somatic mutation lists and RSEM normalized RNA-seq expression data for CESC tumors were obtained from the TCGA at http://gdac.broadinstitute.org/runs/analyses__2016_01_28/data/CESC-TP/20160128/gdac.broadinstitute.org_CESC-TP.CopyNumber_Gistic2.Level_4.2016012800.0.0.tar.gz, http://gdac.broadinstitute.org/runs/analyses__2016_01_28/data/CESC-TP/20160128/gdac.broadinstitute.org_CESC-TP.Mutation_APOBEC.Level_4.2016012800.0.0.tar.gz, and http://gdac.broadinstitute.org/runs/stddata__2016_01_28/data/CESC/20160128/gdac.broadinstitute.org_CESC.Merge_rnaseqv2__illuminahiseq_rnaseqv2__unc_edu__Level_3__RSEM_genes_normalized__data.Level_3.2016012800.0.0.tar.gz, respectively.

### Statistical comparisons of A3B-induced mutation in yeast gene deletion strains

Gene deletion strains with elevated A3B-induced mutation were statistically identified as those with 95% confidence intervals that were non-overlapping compared to wild-type yeast expressing A3B for the median Can^R^ rate determined by the maximum likelihood method in FALCOR. The resulting list of 61 genes were clustered based on related function using string.db (www.sting-db.org) (Szklarczyk et al. 2019) rest to highest confidence linkage and with K-means clustering of 8 nodes. Enriched Component and Process GO categories among these genes were also determined with string.db. Full lists of enriched categories are provided in (Supplemental Tables 7 and 8). Whether specific gene deletions were epistatic, additive, or synergistic with *ung1*Δ in regards to A3B-induced Can^R^ rate was determined by adding the appropriate bounds of the 95% confidence intervals for the single deletion strains to the *ung1*Δ (i.e. upper bound to upper bound and lower bound to lower bound) and comparing these combined bounds to the experimentally determined confidence intervals of the double deletion strains and the *ung1*Δ strain. Epistatic genes contained overlapping experimentally measured confidence intervals with the single *ung1*Δ strain. Additive genes had overlapping measured confidence intervals with those adding the corresponding single deletion strains bounds together. Synergistic genes had higher experimentally determined rates with non-overlapping 95% confidence intervals compared to the estimate calculated by adding bounds from the single deletion strain to the single *ung1*Δ strain. Altered strand biases were identified by the pairwise comparison of the number of C substitutions and G substitutions sequenced *can1* alleles from a specific gene deletion strain expressing A3B to those from A3B expressing wild-type yeast by two-sided Fisher’s Exact test. Altered Rev1-mediated bypass was determined by similar comparison of the ratio of C-to-T and C-to-G substitutions in *can1* from individual gene deletion stains expressing A3B compared pairwise to the same ratio from A3B expressing wild-type yeast by two-sided Fisher’s Exact test. All p-values were corrected for multiple hypothesis testing by the Bonferroni method (Bland and Altman 1995). Hierarchical clustering of gene deletion phenotypes was conducted through the Morpheus webtool (https://software.broadinstitute.org/morpheus/), using average linkage of the one minus Pearson correlation of log2 adjusted values.

### Primary human tumor analyses

Putative APOBEC signature C-to-T and C-to-G mutations were identified among the total list of somatic mutations for whole genome sequenced breast cancers and exome sequenced cervical cancers using the APOBECng module integrated as part of the firehose pipeline (Roberts et al. 2013). Comparisons of APOBEC-induced mutation load between BRCA1/2-proficient and deficient tumors as well as between minus 1 and copy neutral *HPRT1*, *RAD51*, and *RAD51C* cervical cancers was conducted by Mann-Whitney Rank Sum test. TCW to TGW mutations from BRCA1/2-proficient and BRCA1/2-deficient tumors with a statistical enrichment of APOBEC signature mutations (determined by Fisher’s exact test and Benjamini-Hochberg (Benjamini and Hochberg 1995) multiple testing correction as in (Roberts et al. 2013)) were assessed for transcriptional and replicative asymmetry using the asymmetry module in MATLAB (Haradhvala et al. 2016).

## Acknowledgements

We thank Dr. Levi O’Loughlin for initial efforts establishing the methodology underlying our mutagenesis screen and Derek Pouchnik and Mark Wildung for help with Pacific Biosciences SMRT sequencing. We also thank Ellen MacNary for technical help measuring mutation rates and with the genetic screen. This work was supported by National Institutes of Health grants R00ES022633 and R01CA218112 to S.A.R. by NIEHS and NCI, respectively. S.A.R. was also supported by Breast Cancer Research Program Breakthrough Award BC141727 from the Department of Defense. K.L. is supported by NIGMS award R01GM129119.

## Author Contributions

Mating, sporulation, selection of screen strains, and mutation rate measurements were conducted by T.M.M, E.R-R., L.N., A. W., and D. M. T.M.M. and D.M., sequenced the *CAN1* gene to produce mutation spectra. T.M.M, and S.A.R. analyzed mutation rate data and clustered positive screen hits. S.A.R. and N.B. analyzed cancer mutation datasets. T.M.M., K.L., and S.A.R. designed the screen strategy and experiments. T.M.M and S.A.R. wrote the manuscript with editing help from K.L.

