## Supplementary Information for "Genetic modifiers of APOBEC-induced mutagenesis"

SUPPLEMENTAL INFORMATION (List of tables and figures)

Supplemental Table 1: List of all screen hits and candidates

Supplemental Table 2: All mutation rates

Supplemental Table 3: Numeric description of *CAN1* mutation spectra for each genotype

Supplemental Table 4: List of all *CAN1* mutations

Supplemental Table 5: Strain genotypes and details on strain construction

Supplemental Table 6: List and description of oligonucleotides used

Supplemental Table 7: Enrichment component for GO analysis

Supplemental Table 8: Enrichment process for GO analysis

Supplemental Table 9: Hierarchical clustering values

**Supplemental Figure 1: Characteristics of APOBEC-induced mutagenesis in deletion strains with defective HR-directed repair, H3K56ac, and CTF18-RFC.** (A) For yeast strains with deletions known to result in defective homology-directed repair, the A3B-induced $Can^R$ rates for yeast strains with and without in *ung1*Δ, rate of mutations at C and G nucleotides (stand-bias of A3B-induced mutations), and spectra of A3B-induced substitutions (indicative Rev1-dependent TLS usage) are compared. (B) For yeast strains with deletions of genes encoding proteins that modulate H3K56ac, the A3B-induced $Can^R$ rates for yeast strains with and without in *ung1*Δ, rate of mutations at C and G nucleotides (stand-bias of A3B-induced mutations), and spectra of A3B-induced substitutions (indicative Rev1-dependent TLS usage) are compared. (C) For yeast strains with deletions of genes encoding members of the CTF18-RFC complex, the A3B-induced $Can^R$ rates for yeast strains with and without in *ung1*Δ, rate of mutations at C and G nucleotides (stand-bias of A3B-induced mutations), and spectra of A3B-induced substitutions (indicative Rev1-dependent TLS usage) are compared.

### Supplemental Figure 1

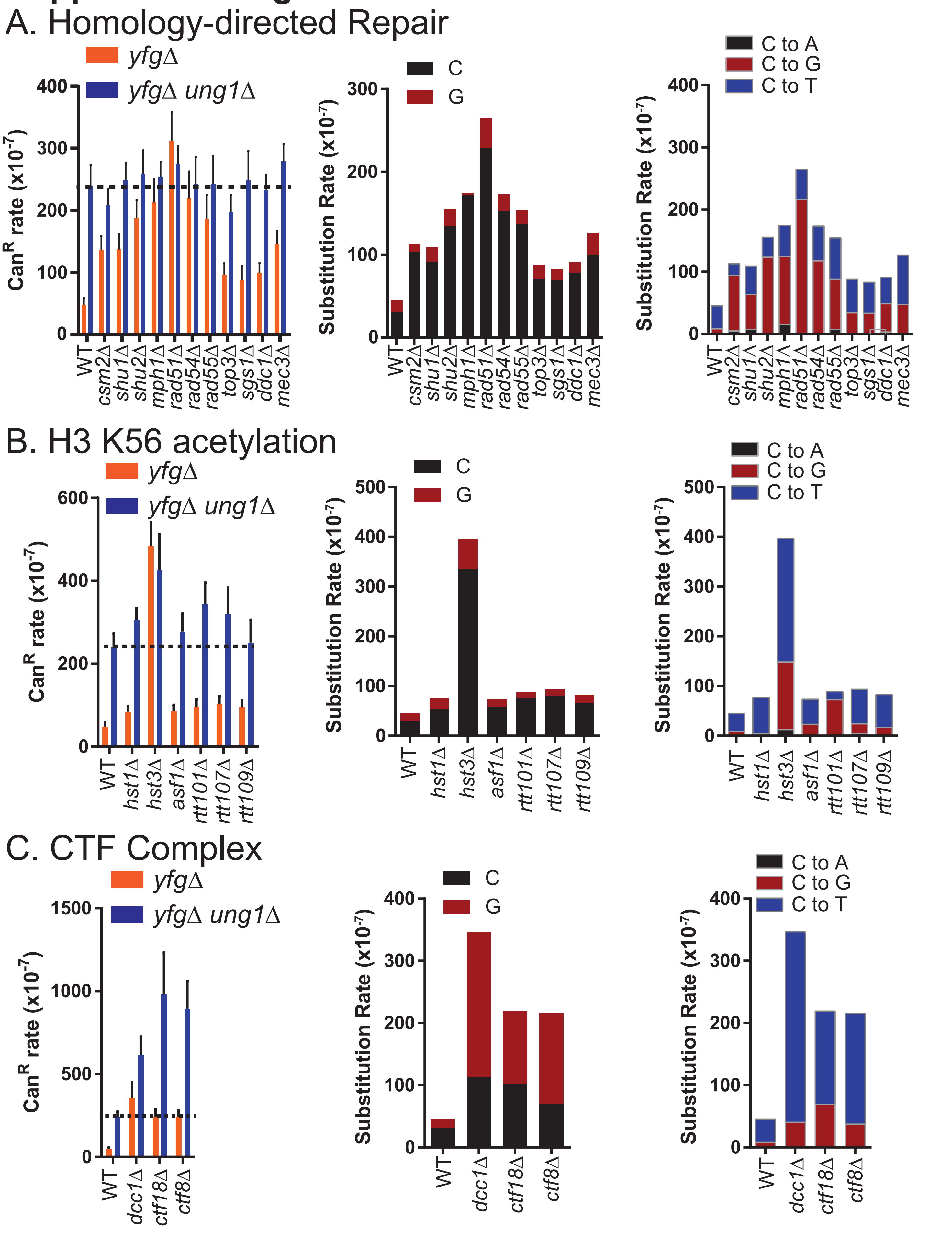
